# Optimizing cardiac organoid culture to enhance maturation, viability, and cardiotoxicity assessments

**DOI:** 10.1101/2025.04.29.651308

**Authors:** Anirudha Harihara, Khashayar Moshksayan, Nima Momtahan, Adela Ben-Yakar, Janet Zoldan

## Abstract

Development of relevant, human induced pluripotent stem cell-derived cardiac organoids is essential to recapitulate myocardium physiology and functionality for assessment of drug-induced toxicity evaluations. However, the optimal conditions for culturing self-aggregating multicellular cardiac organoids are not well-elucidated, particularly the impact of noncardiomyocytes. In this study, we generated cardiac organoids at varying seeding densities to formulate organoids that meet or exceed the biological diffusion limit. We assessed their morphology, gene expression profiles, beating functionality, viability, and mitochondrial activity over time. Our results show that organoid sizes stabilize by seven days of culture, regardless of seeding density. However, organoids seeded with 20,000 cells retained an optimal cardiac signature that promotes cardiac maturity and minimizes fibrotic tendencies, especially when culturing longer than seven days. While all organoid populations maintained their beating functionalities, those seeded with 80,000 cells exhibited greater cell shedding and increased apoptosis at long term culture. In contrast, minimal apoptosis was observed in organoids seeded with 20,000 cells after seven days. Mitochondrial staining further revealed that organoids seeded with 20,000 cells consistently demonstrated higher metabolic activity. Taken together, organoids seeded with 20,000 cells and cultured for seven days yielded the healthiest morphology, transcriptional signature, and viability, while maintaining robust beating kinetics. Importantly, compared to 2D cultures, these optimized organoids demonstrate improved sensitivity to clinically relevant doxorubicin-induced cardiotoxicity, enabling more accurate dose-response evaluations that better reflect therapeutic conditions.

**Impact Statement:** This work highlights key tissue engineering considerations for generating self-assembling cardiac organoids suitable for scalable, high-throughput drug screening and discovery. Understanding the effect of seeding density and culture duration on organoid size and, consequently on gene expression, beating functionalities, apoptosis, and metabolic activity, has broader implications for establishing optimal organoid culture conditions. These insights enable the production of large quantities of cardiac organoids capable of modeling drug-induced toxicity effects on a clinically relevant timescale.

## Introduction

Developing biologically relevant cardiac models is crucial for improving patient-specific drug testing and predicting cardiotoxicity. Between 1950 and 2017, unforeseen myocardial toxicity led to the withdrawal of 61 drugs,^1^ underscoring the limitations of traditional preclinical models. Animal studies often fail to fully replicate human cardiac physiology, while clinical trials frequently lack sufficient genetic diversity.^2^ Advancing *in vitro* human cardiac models offers a transformative solution for predicting patient-specific cardiotoxicity with greater precision.^2^

While primary patient-derived cardiac cells provide an ideal model, their limited viability and invasive collection methods pose challenges.^3^ In contrast, human induced pluripotent stem cells (hiPSCs) offer an abundant, minimally invasive source for generating patient-specific cardiomyocytes.^4^ Traditional 2D cultures provide valuable insights into cellular interactions but fall short in fully capturing the complex cardiac microenvironment.^5^ The next frontier lies in 3D myocardial models, which more accurately replicate cardiac structure and function, thereby improving drug toxicity assessments.^6,7^

Among 3D cardiac microphysiological tissue constructs, hiPSC-derived cardiac organoids present a significant breakthrough. Unlike 2D cultures or other 3D models, organoids self-assemble into microtissues that better mimic the heart’s microenvironment. While engineered cardiac tissues such as Biowire offer functional advantages,^8,9^ they require large cell numbers and involve complex fabrication processes. Similarly, embryoid body-based organoids show potential but suffer from poor control over cell fate, leading to high variability.^10^ Differentiation of micropatterned embryoid bodies can result in ventricular structural formation,^11^ but their stationary nature limits nutrient diffusion, potentially limiting long-term culture.

Multicellular cardiac organoids provide a robust, reproducible model that better recapitulates the myocardium, making them a powerful tool for regenerative medicine and precision therapeutics. Co-culturing cardiomyocytes with fibroblasts enhances the physiological relevance of these models, as fibroblasts contribute essential extracellular matrix components.^12^ However, the influence of fibroblast-cardiomyocyte interactions on organoid necrosis, fibrosis, and cardiac functionality remains poorly understood.

This study investigates how organoid size, which dictates necrotic region formation, impacts cardiac tissue health and function, especially under conditions where nutrient supply may be limited. We investigated two organoid size distributions: one that initially matches the myocardial intercapillary distance of 400 µm,^13^ and another twice its size. Our findings suggest that organoids formed from hiPSC-derived cardiomyocytes and fibroblasts, seeded with 20,000 cells and cultured for seven days, have a stable size, exhibit an optimal cardiac gene expression signature, demonstrate the lowest apoptotic activity, and preserve cardiac functionality. These results provide insights into the relationship between organoid size and cardiac health, offering a more effective model for studying drug-induced cardiotoxicity with fewer cells and improved experimental ease. Finally, we show that this model yields more clinically relevant cardiotoxicity results when exposed to doxorubicin (DOX), reinforcing the potential of hiPSC-derived cardiac organoids as a superior platform for cardiotoxicity assessment during preclinical drug screening.

## Methods

### hiPSC culture and cardiac differentiation for downstream organoid formation

hiPSCs (WTC11) transfected with CMV-GCamp2 (a gift from Dr. Bruce Konklin) were differentiated into cardiomyocytes via WNT/*β*-catenin-pathway modulation,^14^ as we previously described^15^ (Supplementary Fig. S1). Spontaneously beating cardiomyocytes typically emerged 15 days after differentiation initiation, at which point cells were prepared for organoid formation. Briefly, cells were resuspended in RPMI 1640 (Hyclone^™^) supplemented with B27 (Gibco^™^) containing insulin (RB^+^) and 10 µM Rock Inhibitor (Y-27632, RI) and seeded into individual round bottom, ultra-low attachment 96-well plates (CoStar®, Corning) at seeding densities of 20,000 or 80,000 cells per well to generate two distinct organoid size distributions, termed 20KOrg and 80KOrg, respectively. To facilitate cell aggregation, plates were spun down at 300g for 5 minutes. After 48 hours, media was replaced with RB^+^ without RI and subsequently replenished every 48 hours with RB^+^. Parallel monolayer cultures were maintained in multiwell plates to serve as 2D controls for downstream experiments.

### Analysis of organoid size distribution across culture time

To measure the degree of organoid compaction with time, individual organoids were imaged across 14 days, and relative changes in the area of a specified organoid were measured. Brightfield images were obtained using an inverted EVOS FL microscope with 4× (NA 0.13) and 10× (NA 0.3) objectives. We developed a custom plug-in in FIJI/ImageJ to automatically measure the organoids’ sizes. Briefly, a Gaussian filter was applied to the image to eliminate the impact of unaggregated cellular debris. Foreground organoid segmentation was achieved by thresholding based on moment-preservation described by Tsai *et al*.^16^ We report the distributions using the organoids’ effective diameters (*d*_*eff*_), calculated as:

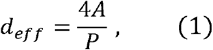

where A and P correspond to the organoid’s area and perimeter, respectively.

### Flow Cytometry

Prior to organoid formation, harvested cells were characterized using flow cytometry to quantify the cardiac differentiation efficiency. Fixed cells were divided into separate suspension to be individually exposed to primary antibodies anti-cTnT (mouse, Invitrogen^™^, 1:300), anti-vimentin (mouse, Invitrogen^™^, 1:100), and anti-CD31 (mouse, Abcam, 1:200) and subsequently exposed to the anti-mouse Alexa488® secondary antibodies (rabbit, Abcam, 1:1000). Cells were then characterized with an Accuri® C6 Flow Cytometer. All analyses of flow cytometry experiments were performed on FlowJo (BD Biosciences).

### Immunostaining of whole organoids

Fixed organoids were exposed to primary antibodies anti-cTnT (rabbit, 1:500) and anti-vimentin (mouse, 1:100) or cleaved caspase 3 (CC3), rabbit, 1:100) diluted in blocking buffer overnight at 4°C. The following secondary antibodies were used: AlexaFluor® 488 (donkey, Invitrogen^TM,^ 1:1000) for anti-vimentin, CF®488 for anti-CC3 (goat, Sigma Aldrich, 1:1000), and AlexaFluor® 594 (Invitrogen^TM,^ 1:1000) for anti-cTnT. For nuclear staining, we used 4’,6’-dimidino-2-phenylinodole (DAPI, Invitrogen^™^, 1:10,000). Organoids were imaged on a confocal microscope (Leica TS SP8) with a 10× (NA 0.3) objective in flat, glass bottom chamber slides (Ibidi).

### Quantitative reverse transcription-polymerase chain reaction (qPCR)

RNA was extracted from 3×10^6^ cells from 2D controls, 150 organoids seeded with 20,000 cells, and 38 organoids seeded with 80,000 cells on Days 2, 7, and 14 of culture using an RNeasy Mini Kit (Qiagen), per the manufacturer’s protocol. Reverse transcription was performed using a High-Capacity cDNA Reverse Transcription Kit (Applied Biosystems). qPCR was performed on a StepOnePlus Real-Time PCR system (Applied Biosystems) using PowerUp SYBR Green (ThermoFisher) and PrimeTime and Prime^™^ qPCR primers (Integrated DNA Technologies, Bio-Rad), listed in Supplementary Table S1. TATA-binding protein (TBP) was used as the endogenous control, and the number of mRNA copies generated on Day 0, at the time of organoid formation, was used as the reference control sample for all experimental timepoints for 2D and 3D samples.

### Beating functionality analysis via calcium transients imaging

GCaMP signaling was imaged over 10 seconds at 100 frames per second on an IX73 Inverted Microscope (10×, 0.3 NA objective, Olympus) at 37°C and 5% CO_2_. Image acquisition was performed using an ORCA-Flash 4.0 V2 Digital CMOS camera C11440-22CU (Hamamatsu Photonics, K.K.).

Image sequences were processed and analyzed using an in-house MATLAB (MathWorks) code to measure beating areas and extract organoid beating waveforms to analyze beating rates, homogeneity, and beating kinetics (Supplementary Fig. S2)

To assess beating homogeneity, i.e., the number of independent calcium transient waveforms the organoids exhibited was measured. We first counted the mean number of independent waveforms within each organoid and then applied a synchrony deviation score to quantify the degree of homogeneity shown by:

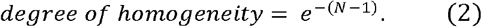

This score ranges between 0 and 1, where N corresponds to the number of detected independent waveforms. A score of 1 indicates perfect homogeneity, with all beating regions exhibiting synchronized calcium transients. A lower score indicates low beating homogeneity, with distinct regions exhibiting different, independent calcium transient waveforms.

Beating rates were determined by counting the number of global maxima in a ten-second recording and multiplying by six to convert to beats per minute (BPM). To analyze the beating kinetics of the organoid waveforms, we measured the full width half maximum (FWHM) and the rise and decay times. The rise time was defined as the interval from the local minimum (baseline signal) to the subsequent signal peak. The decay time was defined as the interval from the peak until the signal decayed to 25% of its peak value.

### Analysis of cleaved caspase 3 staining

Maximum z-projections of confocal image z-stacks stained with DAPI and cleaved caspase-3 (CC3) were generated to identify apoptotic cells. CC3 is a well-established marker of apoptosis, as it indicates activation of the caspase cascade leading to programmed cell death. Apoptotic cells were defined as those stained positively for both DAPI and CC3. The fraction of apoptotic cells was measured as the number of DAPI^+^ and CC3^+^ cells divided by the total number of DAPI^+^ cells.

### Mitochondrial analysis

Following manufacturer instruction, we stained live organoids with MitoTracker^™^ (500nM, ThermoFisher) on Day 7 and again on Day 14 of organoid culture to monitor mitochondrial activity of the organoids as a function of both initial seeding density and time in culture. Stained organoids were imaged using wide-field fluorescence microscopy using an IX73 Inverted Microscope (10×, 0.3 NA objective, Olympus) and were analyzed by evaluating the organoids’ mean integrated intensity.

### Doxorubicin response evaluation of 2D cultures versus 3D organoids

2D and 3D cardiac models were exposed to DOX concentrations of 0.1, 0.3, 0.5, 1, and 10 µM (in 0.1% (v/v) DMSO), along with a vehicle control. Beating was evaluated at 24- and 48-hours post-drug exposure by imaging calcium transients. All imaging was performed using a Cytation 3 Biotek plate reader maintained at 37°C. Beating regions were manually identified and programmed into the plate reader to image the same beating regions at subsequent time points. For beating analysis, individual beating regions of interest (ROI) in both 2D cultures and 3D organoids were tracked over time. Changes in beating rate (ΔBR) relative to their baseline, pre-treatment (pre-tx), values were calculated using:

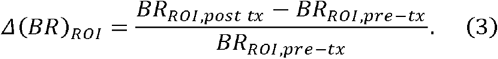

After 48 hours of treatment, both organoids and 2D controls were stained for live and dead cells using Calcein AM (Invitrogen^™^) and Ethidium Homodimer-1 (EthD-1, Invitrogen^™^), respectively. The dyes were administered at 2 µM in RB^+^ medium, per manufacturer’s instructions, and incubated for 30 minutes at 37°C and 5% CO_2_. Live/dead imaging was performed on the Cytation 3 Biotek plate reader using z-stacks acquisition for both 2D and 3D cultures. For viability analysis in 2D cultures, we used maximum intensity projections to integrate the green (live, *S*_*green*_) and red (dead, *S*_*red*_) signals, from which we computed their mean values for each organoid. The viability ratio (VR) is defined according to:

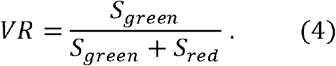

To analyze organoid viability on the captured z-stacks, we created a mask using the brightfield image of the organoid to define the organoid’s boundary. This mask was then applied to the maximum intensity projection images of the corresponding fluorescence images to determine the mean intensities, *S*_*green*_ and *S*_*red*_, identical to the protocol employed for analysis of the 2D controls. The VR of each organoid was calculated according to Eq. 4 and normalized to the average VR of vehicle-treated organoids to determine relative viability.

### Statistical analysis

Statistical analyses were performed using MATLAB and OriginPro (OriginLab Corporation, version 9.9.0.225), including all curve fittings for the dose-response assessments and IC_50_ calculations. We calculated the *p*-values for group comparisons using t-tests, Levene tests, or two-way ANOVA tests where *p* < 0.05 (*), *p* < 0.005 (**), and *p* < 0.0005 (***) were considered significant. All data points in data are presented as mean ± standard error of the mean (SEM) unless otherwise stated.

## Experiment

### Organoid formation and size stabilization results in prolonged maintenance of organoid morphology

We sought to determine the size dimension at which cardiac organoids stabilize over time. We hypothesized that organoids formed solely via gravitational sedimentation would exhibit a heterogeneous size distribution despite the round-bottom of the wells because their relatively wide curvature at the base. To address this hypothesis, we introduced a two-minute, 200g centrifugation step immediately after seeding cells into the wells. This centrifugation produced an adequate centripetal force to facilitate cell aggregation into a single organoid, leading to organoids sizes that were more reproducible and directly proportional to seeding density (Supplementary Fig. S3). Without centrifugation, cells typically formed one main organoid, accompanied by adjacent smaller cell clusters.

Next, we studied the impact of cell seeding density on organoid size. We hypothesized that the presence of fibroblasts would lead to significant organoid compaction, which would depend on the initial cell seeding density used. We tested two seeding densities to generate 20KOrg and 80KOrg (Figs. 1a, b). Over 14 days, the sizes of both types of organoids varied significantly with 80KOrgs exhibiting greater variation. Two days after organoid formation, the average effective diameters of the 20KOrg and 80KOrg were 510.6±5.3 and 774.8±13.5 µm, respectively. By day seven, the organoids compacted, stabilizing to average diameters of 421.5±4.9 and 590.7±6.3 µm for 20KOrg and 80KOrg, respectively. After another week in culture, further compaction occurred, with final average effective diameters of 409.4±6.6 and 488.4±8.0 µm (Fig. 1b).

**Figure 1.**
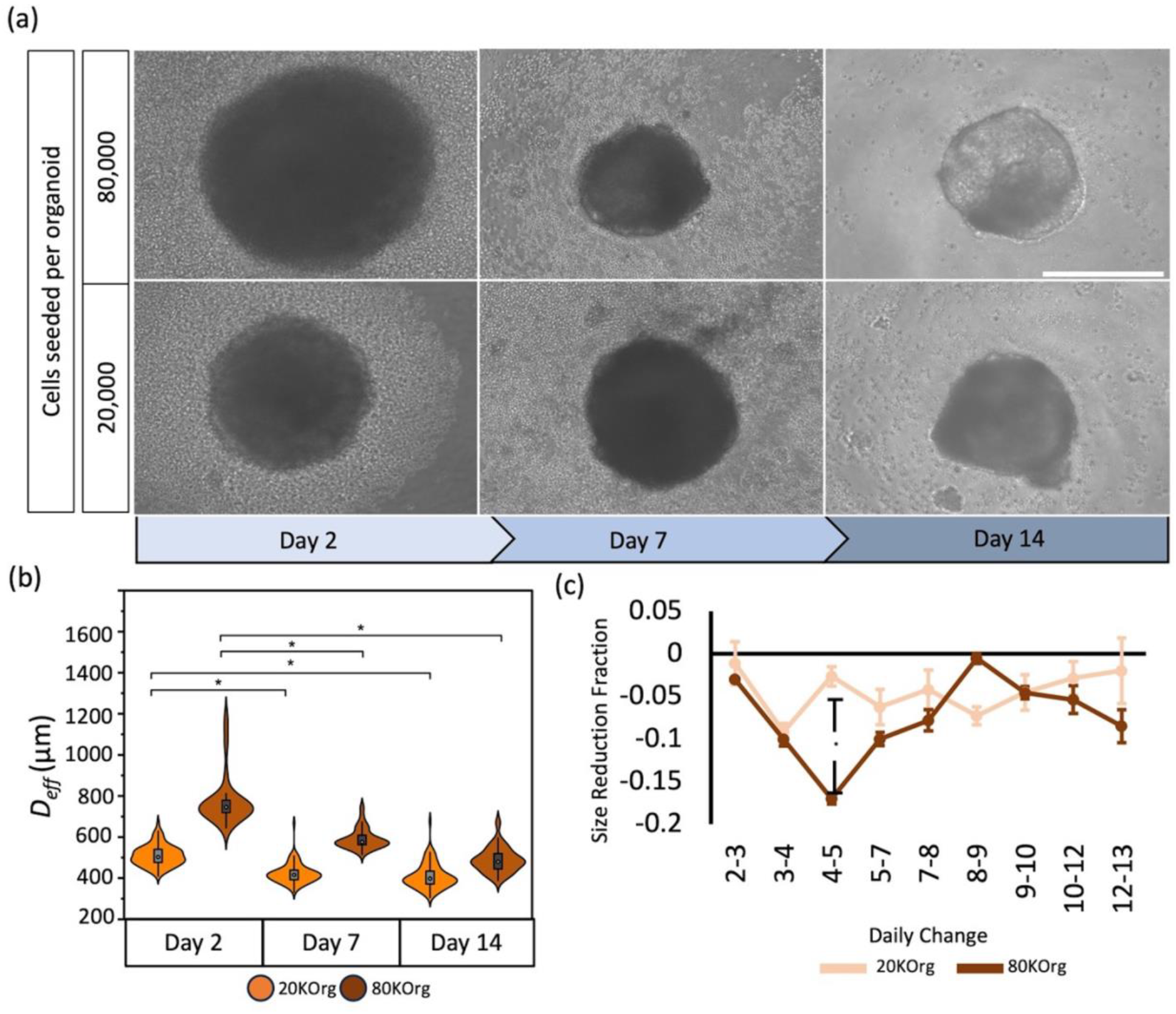
Characterization of organoid morphology and size distributions throughout 14 days of culture. (**a**) Representative images of 20KOrg *(bottom)* and 80KOrg *(top)*. Scale bar is 400 µm. (**b**) Violin plots of effective diameters measured from cardiac organoids measured at three key time points: 2-, 7-, and 14-days post organoid formation (n_20kOrg, day2_=178, n_80kOrg, day2_=112, n_20kOrg, day7_=214, n_80kOrg, day7_=156, n_20kOrg, day14_=161, n_80kOrg, day14_=142). (**c**) Daily imaging of organoids reveals that maximum compaction occurs on Day 3 of organoid culture for 20KOrg. However, 80KOrg undergo maximum compaction at Day 4, after which both conditions are statistically similar during subsequent days of culture.

To determine whether changes in organoid size are solely due to compaction or also influenced by cell shedding, we counted the number of DAPI-stained nuclei within the organoid imaging volume. We observed a significant reduction in nuclei density at Days 7 and 14 for both seeding densities (Supplementary Fig. S4). 20KOrg lost approximately 47.7% of their cells by Day 7 and an additional 65.6% by Day 14. In comparison, 80KOrg showed a 43.4% reduction in cell number by Day 7 and an additional 49.3% decrease by Day 14. After seven days in suspension culture, organoids seeded at higher cell densities showed reduced cell shedding but reached maximum compaction, resulting in no significant size differences compared to those seeded at the lower density (Fig. 1b, c). While organoid size distributions stabilized by Day 7, it was necessary to evaluate their cardiac signatures.

### Assessment of cardiac signature in organoids

We first characterized the composition of the initial cell populations used for organoid formation. Based on our experience, we anticipated that cardiac differentiation via WNT/*β*-catenin pathway modulation would yield a heterogenous cell population comprised of cardiomyocytes, fibroblasts, and endothelial cells.^17,18^ Immunostaining and flow cytometry confirmed the presence of both cardiomyocytes (22.31±6.38 %) and fibroblasts (76.40±5.88 %) with a minor fraction of endothelial cells (1.29±1.01%) (Supplementary Fig. S5), indicating that the cell populations used to form cardiac organoids were predominantly cardiomyocytes and fibroblasts. The low percentage of endothelial cells suggests that vascularization within the organoids was unlikely, predisposing larger organoids, especially those exceeding the diffusion limit, to develop apoptotic and necrotic areas. Immunostained organoids on Day 14 confirmed that the organoids retained their composition of cTnT-positive cardiomyocytes and vimentin-positive fibroblasts (Fig. 2a).

**Figure 2.**
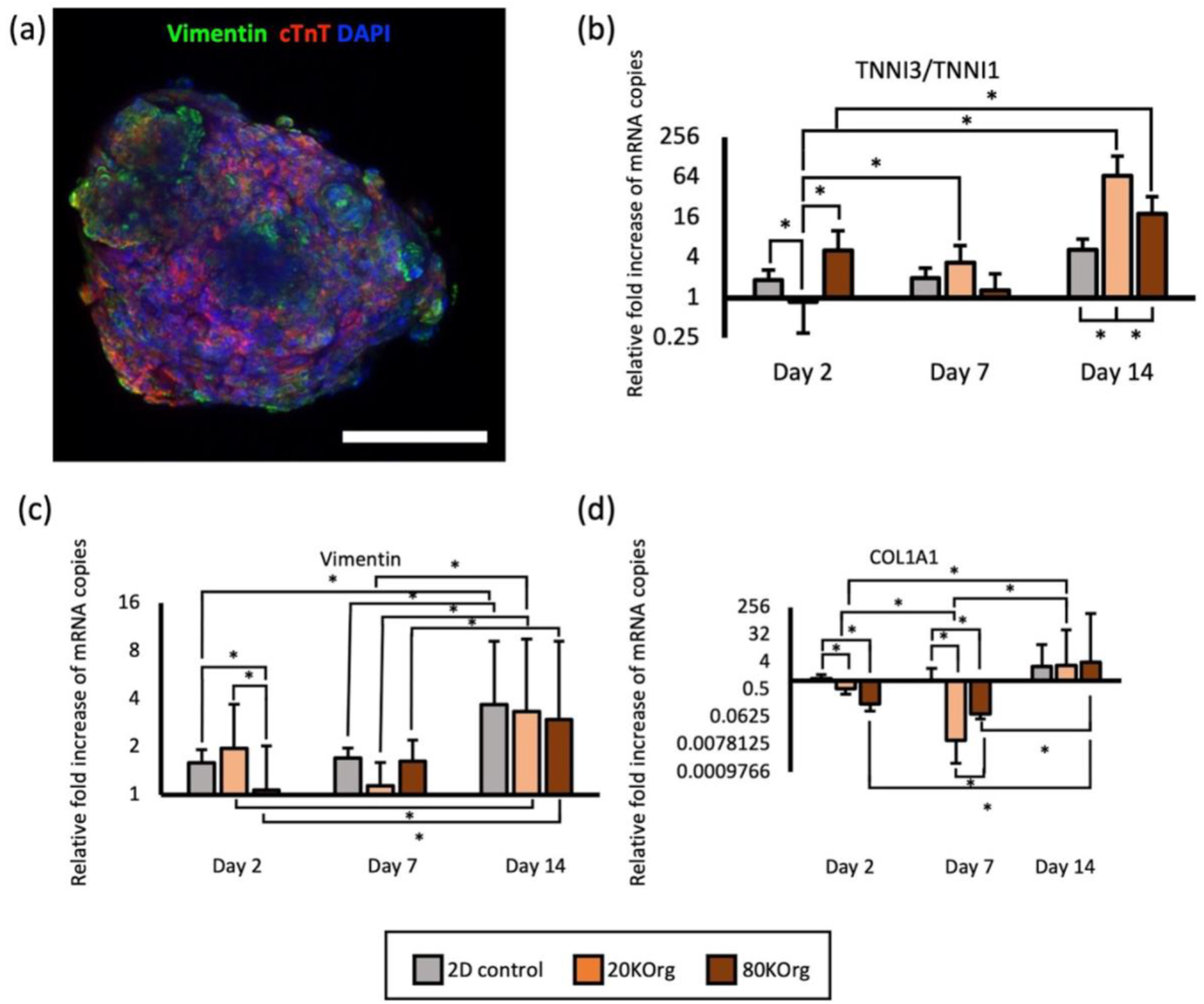
Determination of cardiac maturation and fibrotic markers in organoid population. (**a**) Z-projection of 20KOrg stained for troponin-positive cardiomyocytes (anti-cTnT, red) and fibroblasts (anti-Vimentin, green) on Day 14. DAPI-stained cell nuclei shown in blue; Images performed using confocal microscopy. Scale bar is 200 µm. (**b-d**) qPCR plots showing relative fold increase of mRNA copies in 2D controls (n=3 wells) 20KOrg (n=468) and 80KOrg (n=108) for TNN3/TNNI1 (**b**), vimentin (**c**), and COLA1A1 (**d**). Unless otherwise indicated, all bars indicate statistical significance of *p* < 0.05. Statistical comparison of gene expression performed using two-way ANOVA testing.

To further understand the dynamics of organoid development, we analyzed gene expression profiles across seeding densities over time. We studied the relative expression levels of TNNI1 and TNNI3 to evaluate cardiomyocyte maturity, vimentin for fibroblasts, and collagen type I (COL1A1) for activated fibroblasts (Figs. 2b-d). As expected, relative TNNI3/TNNI1 expression levels, indicative of cardiac maturation, increased over time in all culture conditions, and even more so for 3D organoid models, aside from 80KOrg cultured for 7 days. On Day 2, 80KOrg exhibited a six-fold upregulation of TNNI3/TNNI1 compared to 20KOrg. However, with longer culture time, this trend reversed, with 20KOrg showing a 2.63-fold and 18-fold increase in TNNI3/TNNI1 expression on Days 7 and 14, respectively, compared to 80KOrg. Next, we assessed if tendencies towards fibrosis were also enhanced over time. We observed a general increase in vimentin expression over time, with a transient decrease on Day 7 in 20KOrg, followed by a three-fold upregulation across all conditions by Day 14. COL1A1 expression was significantly downregulated in 20KOrg (91-fold) and 80KOrg (12-fold) on Day 7 compared to 2D controls, demonstrating increased cellular health in younger organoid models, that eventually increased in all models by Day 14.

To assess optimal cardiac health across conditions based on all gene expression patterns, we performed unsupervised hierarchical clustering on normalized gene expression (ΔΔCT) values in MATLAB. This analysis identified the optimal culture condition as 20KOrg cultured for seven days, which maintained high TNNI3/TNNI1 expression while minimizing fibrotic markers (COL1A1, vimentin) (Supplementary Fig. S6). To validate this, we further analyzed their beating behavior, apoptosis, and metabolic activity.

### Beating assessment of cardiac organoids for determination of optimal organoid health

We characterized calcium transients in spontaneously beating organoids on Days 7 and 14. Figures 3a and 3b show identified beating regions and corresponding waveforms for 20KOrg and 80KOrg, respectively. Beating rates and associated parameters did not differ significantly between seeding densities or over time. 20KOrg exhibited beating rates of 79±8 bpm on Day 7 and 60±6 bpm on Day 14, while 80KOrg had rates of 57±11 bpm and 56±11 bpm, respectively (Fig. 3c).

**Figure 3.**
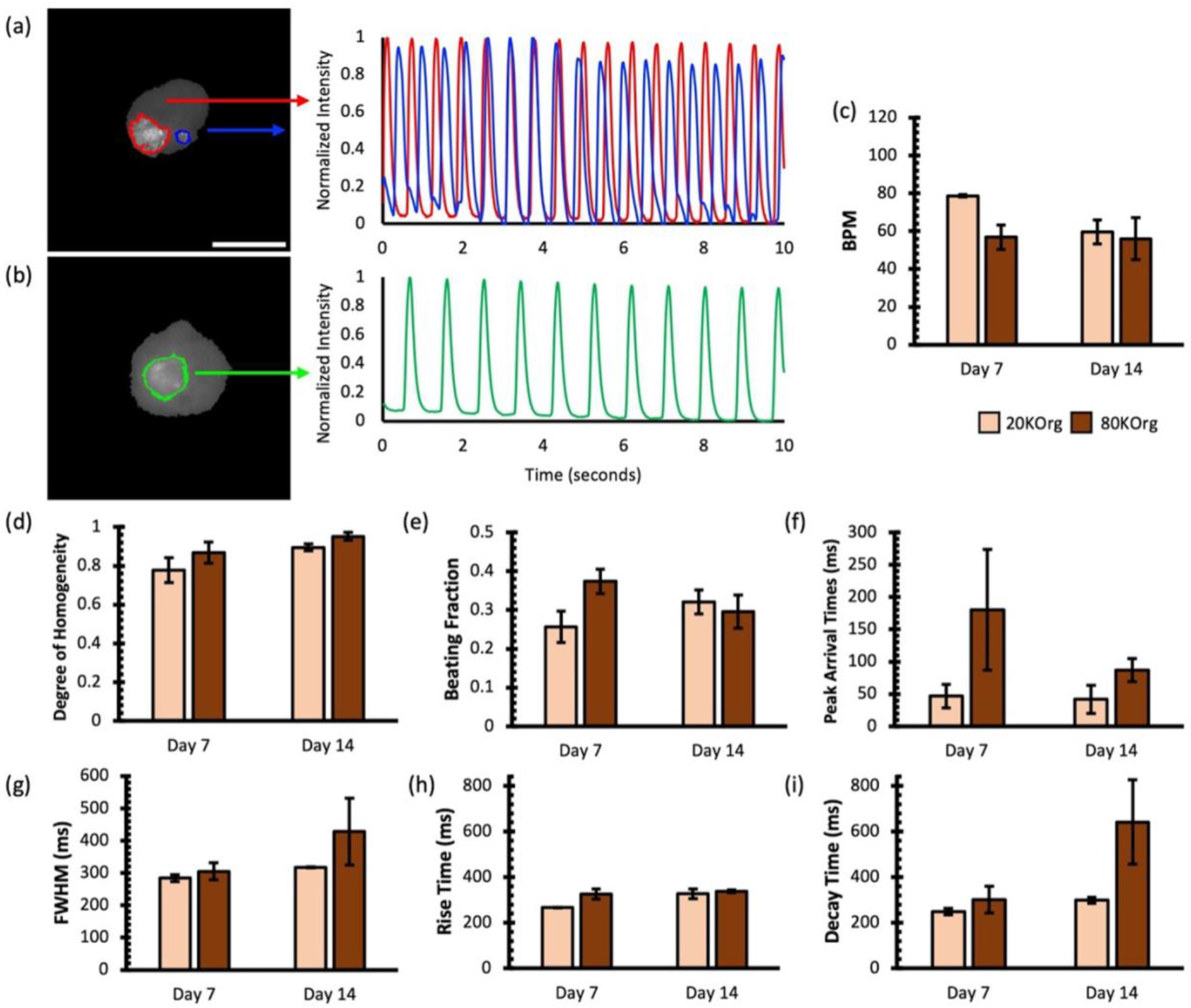
Assessment of organoid beating functionality. **(a)** Segmented ROIs of beating regions within 20KOrg on Day 7 and **(b)** 80KOrg on Day 7 along with their corresponding beating waveforms derived from the standard deviation projection of pixel intensity changes over the entire image sequence. Scale bar is 400 µm. **(c)** Beats per minute of organoid culture groups on Days 7 and 14. **(d)** Degree of homogeneity, **(e)** Beating fraction of organoids determined by regions exhibiting fluctuating calcium transients over time at a specified focal plane. **(f)** Peak arrival times of calcium propagation across organoid beating regions. **(g)** Full width half maximal characterizations to approximate beating durations of organoid populations. **(h)** Rise times of organoid waveforms defined as the duration required to reach the maximum calcium signal intensity relative to the baseline signal (adjacent prior local minima). **(i)** Decay times of organoids waveforms defined as the duration from the maximum calcium signal intensity to 25% of its peak value (n_20KOrg, Days 7 & 14_=12, n_80KOrg, Day 7_=23, n_80KOrg, Day14_=20). Statistical analysis of beating parameters was performed using two-way ANOVA followed by Tukey’s post hoc test for multiple comparisons.

Waveform homogeneity improved over time, with values of 0.67±0.08 (Day 7) and 0.83±0.02 (Day 14) for 20KOrg, and 0.85±0.06 (Day 7) and 0.92±0.03 (Day 14) for 80KOrg (Fig. 3d). Projected beating fractions were 0.25±0.04 and 0.32±0.03 on Days 7 and 14 for 20KOrg, and 0.37±0.03 and 0.30±0.04 for 80Korg (Fig. 3e), with no significant differences likely due to organoid size stabilization by Day 7.

Peak arrival times were highly variable but averaged 46.8±18.2 ms (20KOrg) and 180.3±93.7 ms (80KOrg) on Day 7 (Fig. 3f). 80KOrg displayed longer FWHM values, rise times, and decay times, particularly on Day 14, correlating with their lower beating rates (Figs. 3g–j).

Despite no significant differences in overall beating kinetics, 20KOrg demonstrated more stable synchronicity and peak arrival times, suggesting better recapitulation of healthy cardiac function. However, the persistence of beating across all conditions indicates their potential for modeling different myocardial states, including mature, immature, and fibrotic phenotypes for cardiotoxicity studies. To further define the healthiest organoid model, we examined apoptotic patterns.

### Apoptotic behavior observed in organoids across seeding densities and culture periods

We hypothesized that the lack of vascular networks in the organoids would result in inadequate passive diffusion of nutrients into the organoids’ core, resulting in higher apoptotic activity, particularly in conditions with increased proclivity towards fibrosis. As caspase 3 cleavage (CC3) has been reported as an initiator of apoptosis in various cell types,^19^ including cardiac,^20^ we immunostained organoids to identify CC3 expression and assessed the degree of apoptosis across time and seeding density.

On Day 2, CC3^+^ fractions were higher in 20KOrg (0.240±0.062) than in 80KOrg (0.190±0.020; Figs. 4a, b). However, by Day 7, apoptosis significantly decreased to 0.083±0.013 in the 20KOrg, while only slightly dropping to 0.179±0.007 in the 80KOrg. By Day 14, apoptosis increased in both groups, reaching 0.158±0.029 and 0.275±0.040, respectively, though 20KOrg maintained significantly lower apoptotic levels, indicating greater resilience over time. Interestingly, while 20KOrg initially had higher CC3 expression on Day 2, it declined significantly by Days 7 and 14, whereas apoptosis in the 80KOrg remained high and increased by Day 14. Comparing these trends with size distribution and gene expression data, we found that although organoid size stabilized by Day 7 due to compaction and cell shedding, larger organoids (80KOrg) exhibited more fibrotic tendencies, likely leading to greater apoptosis. This supports our hypothesis that smaller organoids (20KOrg), which better match intracapillary distances in cardiac tissue, exhibit reduced apoptosis. Next, we examined whether this apoptotic activity correlated with metabolic differences.

**Figure 4.**
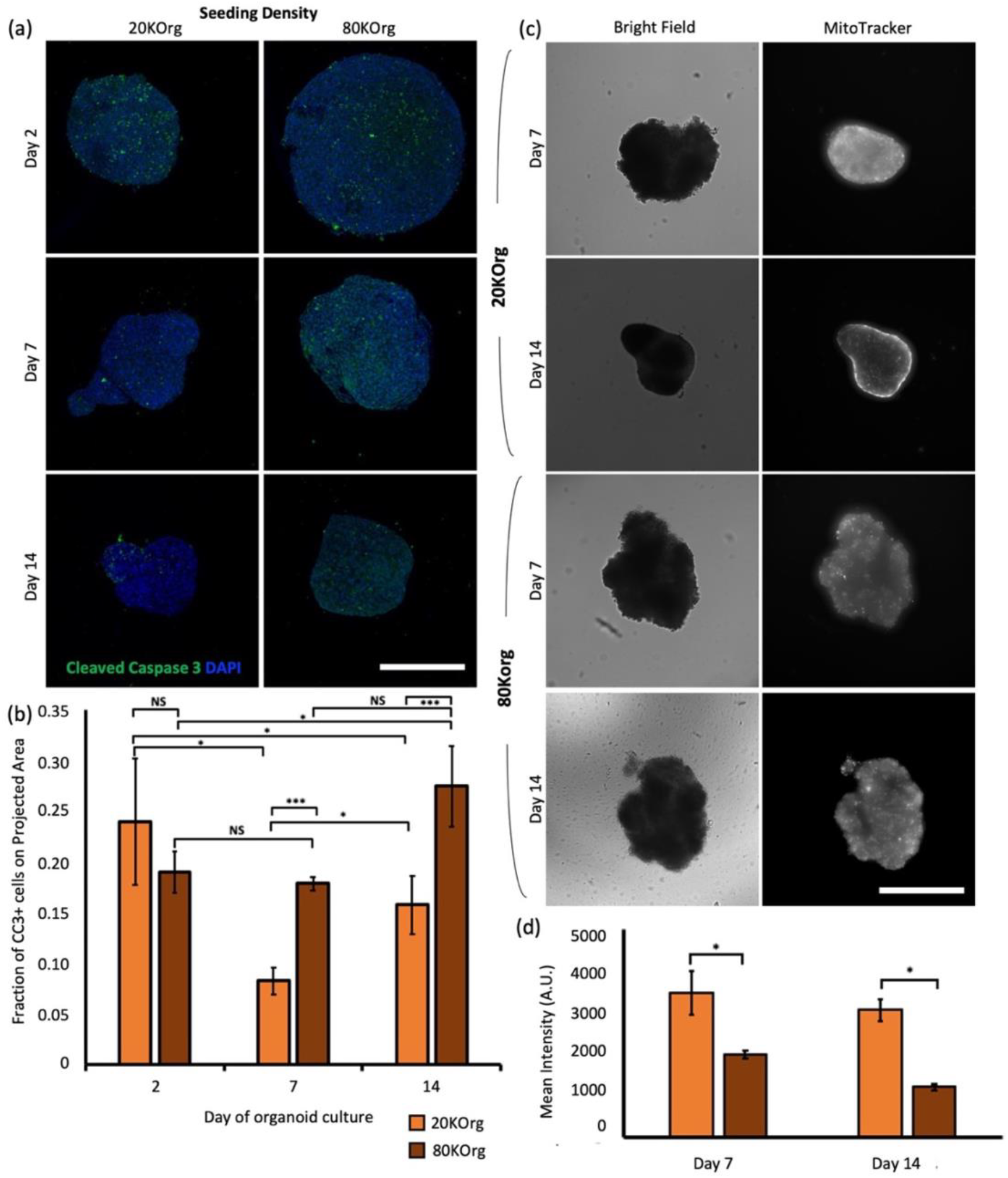
Determination of organoid proneness as a function of size and metabolic activity. **(a)** Panel of MAX z-projections of image stacks captured at key time points for organoids seeded at the two densities, stained for cleaved caspase 3 (CC3, green) and DAPI (blue) to identify necrotic and apoptotic cells obtained using confocal microscopy. Scalebar is 400 µm. **(b)** CC3*+* necrotic areas in z-projections reveal significantly less apoptosis and necrosis in 20KOrgs. These organoids exhibit sufficient shedding of necrotic regions by Day 7, coinciding with size stabilization. Additional cell death is observed upon Day 7 of culture, corresponding to concurrent onset of organoid fibrosis (n_20KOrg, day2_=4, n_80KOrg, day2_=9, n_20KOrg, day7_=4, n_80KOrg, day7_=5, n_20KOrg, day14_=7, n_80KOrg, day14_=5). **(c)** MitoTracker*TM* staining at Day 7 in 20KOrg (left) and 80KOrg (right). Scalebar is 400 µm. **(d)** Quantification of MitoTracker*TM*-expression shows enhanced mitochondrial activity in 20KOrg, indicating a healthier organoid model for downstream experiments. Disparities between seeding densities persist without significant changes over time (n=7 per condition).

### Mitochondrial activity differences observed between the different seeding densities of cardiac organoids

We hypothesized that the higher fibrosis and apoptosis observed in the 80KOrg could affect their metabolic activity. To assess this, we used MitoTracker™ to monitor mitochondrial activity from Day 7 to Day 14 (Fig. 4c). 20KOrg displayed significantly higher mitochondrial intensity than 80KOrg (Fig. 4d). Combined with increased vimentin and COL1A1 expression, this suggests that larger organoids develop fibrotic and apoptotic regions over time, leading to reduced metabolic activity.

Despite both groups showing increased fibrosis and apoptosis by Day 14, mitochondrial activity remained stable within each seeding density. We attribute this to adequate nutrient diffusion sustaining peripheral cells’ metabolic activity. Given their higher metabolic activity, we next evaluated whether 20KOrg could effectively model drug-induced toxicity.

### Organoids better predict cardiotoxicity over 2D models

Based on our findings, we selected 20KOrg to assess their potential as a 3D drug screening model, specifically for testing cardiotoxic drugs like doxorubicin (DOX). Cardiac organoids and 2D cultures were treated with DOX (0.1–10 µM, plus a vehicle control) for 48 hours. Beating was recorded at 24 and 48 hours, while viability was assessed after 48 hours (Figs. 5a, b). Dose-response analysis showed a higher IC_50_ for organoids compared to 2D cultures (Fig. 5c), indicating greater resistance to DOX. Similarly, the change in beating rate (ΔBR) after 24 hours of exposure yielded a higher IC_50_ in organoids (IC_50_=1.48±0.38 µM) than in 2D cultures (IC_50_=0.76±0.0.02 µM), as shown in Figure 5d. By 48 hours, organoids maintained a significantly higher IC_50_ of 0.99±0.39 µM, compared to the measured IC_50_ of 0.47±0.03 µM in 2D cultures (Fig. 5e). Given that reported plasma toxicity levels of DOX range from 2.79–11 µM,^21,22^ these results suggest that 3D cardiac organoids may more accurately model cardiotoxicity than traditional 2D cultures. Furthermore, these organoids exhibit performance comparable to those reported in the literature, particularly in terms of cardiac functionality. Specifically, when Lee *et al*. cultured hiPSC-derived heart organoids on a similar timescale, they report an IC_50_ of 1.23 µM.^23^ Others have shown IC_50_ values on an order similar to our findings. For example, Richards *et. al*. reported an IC_50_ value of 0.41 µM for organoids exposed to DOX for 48 hours, reporting on the fractional area change of cardiac organoids during beating.^24^

**Figure 5.**
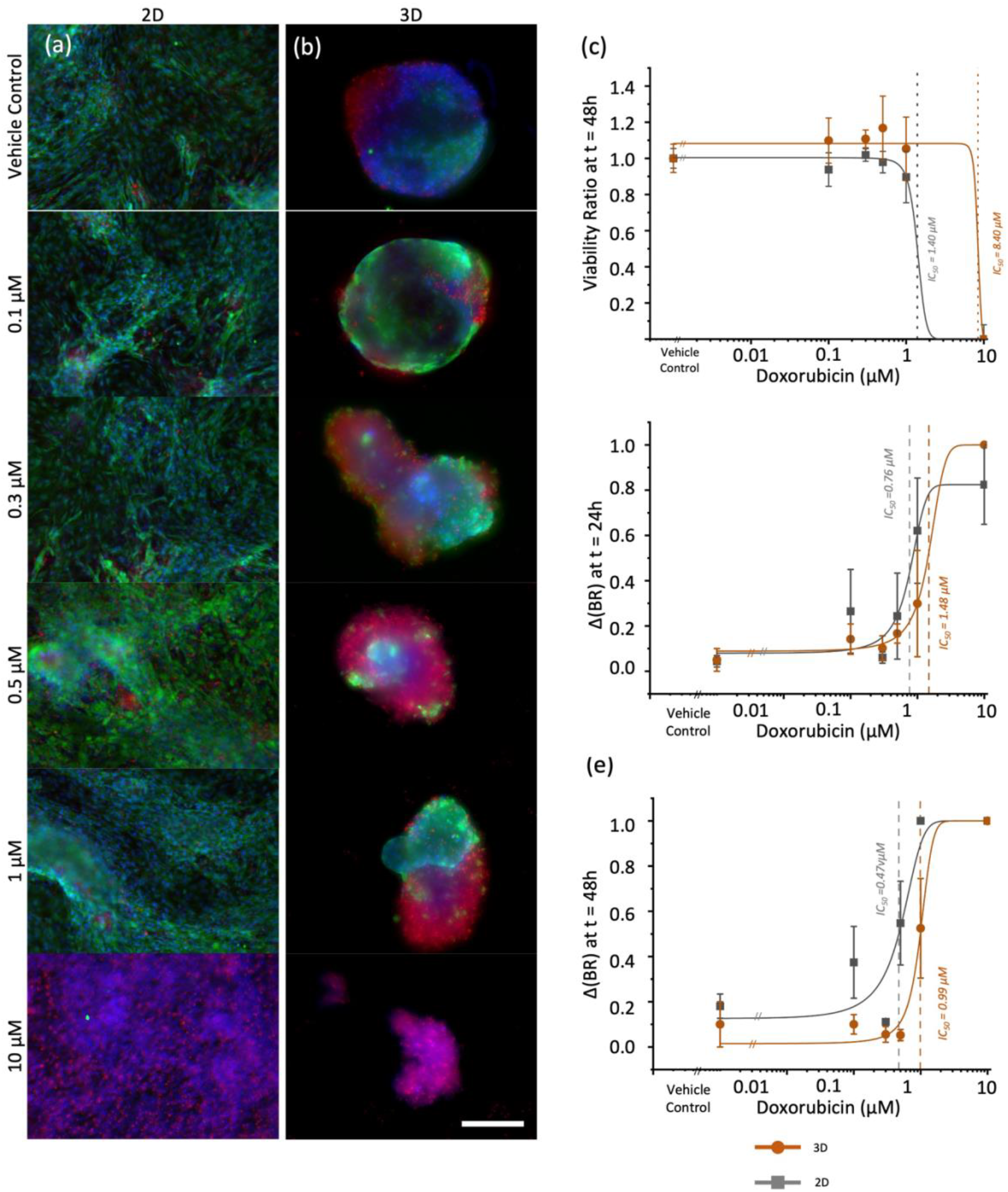
Preliminary drug response studies showcase trend towards more clinically relevant inhibitory concentrations for cardiac organoids. (**a**) Representative images of 2D plated cells stained for live cells (Calcein-AM, green), dead cells (Ethidium Homodimer, red) cells and their nuclei (Hoecsht33342) following 24 hours of doxorubicin exposure. (**b**) Representative images of organoids stained with the same marker following 24 hours of doxorubicin exposure (scale bar for (a) and (b) is 400 µm). (**c-e**) Dose response curve comparing doxorubicin-induced toxicity profiles in 2D and 3D cardiac models with respect to viability ratio (**c**), ΔBR at 24-hours (**d**), and ΔBR at 48-hours (**e**); (n*2D (vehicle)*=3, n*2D (0*.*1µM, 0*.*5µM, 1µM)*=6, n*2D* (*0*.*3µM)*=5, n*2D (10µM)*=4, n*3D*=4 per concentration).

## Discussion

Herein, we describe our determination of the optimal self-assembled cardiac organoid candidate for downstream analysis. We propose that culturing 20KOrg with a gentle centrifugation step in round bottom ULA plates can serve as the healthiest model for downstream cardiotoxicity studies. This is evidenced by its most stable size distribution across 14 days with reduced cell shedding, while maintaining native beating rates and parameters. Particularly, when culturing these organoids for 7 days, we report a more optimal cardiac signature with minimal fibrosis levels, low apoptosis levels, and higher mitochondrial activity. Furthermore, their stabilization of beating parameters by 7 days of culture suggests they can be used for drug exposure experiments earlier, thus saving time in the clinic when observing drug-induced responses. This method maintains key parameters necessary for cardiac probing while requiring minimal effort in formulation to generate high volumes of samples. It maintains all key cardiac parameters of interest without introducing hydrogels and labor-intensive introduction of alternate cell types to aid in vascularization for nutrient supply. While sophisticated models have demonstrated recapitulation of *in vivo* cardiac physiology,^8-12^ these models require heavy user manipulation, and their generation is very challenging for higher throughput demands. We previously demonstrated that organoids similar to those reported here, when integrated with our microfluidic platform, OrganoidChip, can enable high-throughput, high-content drug screening.^25, 26^

Here we optimized culture conditions to yield cardiac organoids with the healthiest morphology and functionality. Nevertheless, fibrotic cardiac organoids, such as the 80KOrg cultured for 14 days, can serve as an effective tool to mimic the cardiac environment of patients predisposed to cardiac fibrosis, as seen in chronic heart diseases, to study untargeted responses to administered profibrotic drug classes.^27^ These include Angiotensin II, used for high blood pressure, kidney failure, and cardiac hypertension treatment,^28^ and TNF-*α* inhibitors, used for treatment of rheumatoid arthritis, psoriatic arthritis, and inflammatory bowel diseases.29

Similarly, it is vital to use proper *in vitro* models that recapitulate drug-induced responses in patients who suffer from ischemic reperfusion injury, which leads to significant cell death.^30^ Here, we also propose that 80KOrg could serve as a model to characterize drug responses from ischemic cardiac tissue as they demonstrate higher levels of apoptosis at Days 7 and 14 along with a more fibrotic gene expression profile. Cardiac ischemia results from changes in the myocardium’s biochemical environment. These alterations compromise tissue stability and impair electrical signal propagation, ultimately leading to myocardial infarction and/or arrhythmia. In our studies larger organoids displayed lower but regular beating rates, accompanied by smaller active beating regions. These results suggest their increased load to maintain adequate oxygen perfusion, further mirroring the pathological features of ischemic myocardium.

In the future, we plan to examine the cell composition of the supernatant to accurately quantify what contributes to cell shedding during culture and create a culture environment that minimizes cell death. One possible option is to look at methods that encourage vasculature formation via co-cultures with relevant cells within self-assembled organoids without using hydrogels that require multi-step user intervention for co-culturing. Additionally, in light of the FDA Modernization Act 2.0, which approved the use of organoid data for drug response evaluation over *in vivo* animal testing, we propose that our model may help ease the sample preparation process during preclinical screens for drug candidates.

## Acknowledgements

We gratefully appreciate the support of the National Institute of Health (1R01GM148906 to A.B.), the Harry S. Moss Heart Trust (UTAUS-FA00002510 to J.Z.), and the National Science Foundation (2002652 to J.Z.).

## Author Contributions

A.H.: Investigation, Conceptualization, Visualization, Software – Original Draft. K.M.: Investigation, Formal analysis, Validation. N.M.: Investigation, Validation. A.B. and J.Z.: Conceptualization, Resources, Writing – Review & Editing, Supervision, Funding Acquisition.

## Data Availability

The datasets generated and analyzed in the presented study are available from the corresponding author upon reasonable request.

## References

1. Kocadal K. Drug-Related Cardiovascular Risk: Retrospective Evaluation of Withdrawn Drugs. North Clin Istanb 2018; doi: 10.14744/nci.2018.44977.

2. Menna P, Gonzalez Paz O, Chello M, et al. Anthracycline cardiotoxicity. Expert Opin Drug Saf 2012;11(sup1):S21–S36; doi: 10.1517/14740338.2011.589834.

3. Akins RE, Rockwood D, Robinson KG, et al. Three-Dimensional Culture Alters Primary Cardiac Cell Phenotype. n.d. 4.

4. Park IH, Lerou PH, Zhao R, et al. Generation of human-induced pluripotent stem cells. Nat Protoc 2008;3(7):1180–1186; doi: 10.1038/nprot.2008.92.

5. Nicin L, Wagner JUG, Luxán G, et al. Fibroblast-Mediated Intercellular Crosstalk in the Healthy and Diseased Heart. FEBS Lett 2022;596(5):638–654; doi: 10.1002/1873-3468.14234.

6. Mathur A, Loskill P, Shao K, et al. Human iPSC-based cardiac microphysiological system for drug screening applications. Sci Rep 2015;5; doi: 10.1038/srep08883.

7. Charwat V, Charrez B, Siemons BA, et al. Validating the Arrhythmogenic Potential of High-, Intermediate-, and Low-Risk Drugs in a Human-Induced Pluripotent Stem Cell-Derived Cardiac Microphysiological System. ACS Pharmacol Transl Sci 2022;5(8):652–667; doi: 10.1021/acsptsci.2c00088.

8. Nunes SS, Miklas JW, Liu J, et al. Biowire: A platform for maturation of human pluripotent stem cell-derived cardiomyocytes. Nat Methods 2013;10(8):781–787; doi: 10.1038/nmeth.2524.

9. Zhao Y, Rafatian N, Feric NT, et al. A Platform for Generation of Chamber-Specific Cardiac Tissues and Disease Modeling. Cell 2019;176(4):913–927.e18; doi: 10.1016/j.cell.2018.11.042.

10. Lewis-Israeli YR, Wasserman AH, Gabalski MA, et al. Self-assembling human heart organoids for the modeling of cardiac development and congenital heart disease. Nat Commun 2021;12(1); doi: 10.1038/s41467-021-25329-5.

11. Hoang P, Kowalczewski A, Sun S, et al. Engineering spatial-organized cardiac organoids for developmental toxicity testing. Stem Cell Reports 2021;16(5):1228–1244; doi: 10.1016/j.stemcr.2021.03.013.

12. Beauchamp P, Jackson CB, Ozhathil LC, et al. 3D Co-culture of hiPSC-Derived Cardiomyocytes With Cardiac Fibroblasts Improves Tissue-Like Features of Cardiac Spheroids. Front Mol Biosci 2020;7; doi: 10.3389/fmolb.2020.00014.

13. Auger FA, Gibot L, Lacroix D. The pivotal role of vascularization in tissue engineering. Annu Rev Biomed Eng 2013;15:177–200; doi: 10.1146/annurev-bioeng-071812-152428.

14. Lian X, Zhang J, Azarin SM, et al. Directed cardiomyocyte differentiation from human pluripotent stem cells by modulating Wnt/β-catenin signaling under fully defined conditions. Nat Protoc 2013;8(1):162–175; doi: 10.1038/nprot.2012.150.

15. Kalkunte NG, Delambre TE, Sohn S, et al. Engineering Alignment Has Mixed Effects on Human Induced Pluripotent Stem Cell Differentiated Cardiomyocyte Maturation. n.d.; doi: 10.1101/2022.09.26.509194v1.

16. Tsai W-H. NOTE Moment-Preserving Thresholding: A New Approach. 1985.

17. Kalkunte NG, Delambre TE, Sohn S, et al. Engineering Alignment Has Mixed Effects on Human Induced Pluripotent Stem Cell Differentiated Cardiomyocyte Maturation. n.d.; doi: 10.1101/2022.09.26.509194v1.

18. Dzobo K, Vogelsang M, Parker MI. Wnt/β-Catenin and MEK-ERK Signaling are Required for Fibroblast-Derived Extracellular Matrix-Mediated Endoderm Differentiation of Embryonic Stem Cells. Stem Cell Rev Rep 2015;11(5):761–773; doi: 10.1007/s12015-015-9598-4.

19. Jelínek M, Balušíková K, Schmiedlová M, et al. The role of individual caspases in cell death induction by taxanes in breast cancer cells. Cancer Cell Int 2015;15(1); doi: 10.1186/s12935-015-0155-7.

20. Chandrashekhar Y, Sen S, Anway R, et al. Long-Term caspase inhibition ameliorates apoptosis, reduces myocardial troponin-I cleavage, protects left ventricular function, and attenuates remodeling in rats with myocardial infarction. J Am Coll Cardiol 2004;43(2):295–301; doi: 10.1016/j.jacc.2003.09.026.

21. Chekin F, Myshin V, Ye R, et al. Graphene-modified electrodes for sensing doxorubicin hydrochloride in human plasma. Anal Bioanal Chem 2019;411(8):1509–1516; doi: 10.1007/s00216-019-01611-w.

22. He H, Liu C, Wu Y, et al. A Multiscale Physiologically-Based Pharmacokinetic Model for Doxorubicin to Explore its Mechanisms of Cytotoxicity and Cardiotoxicity in Human Physiological Contexts. Pharm Res 2018;35(9); doi: 10.1007/s11095-018-2456-8.

23. Jang J, Jung H, Jeong J, et al. Modeling doxorubicin-induced-cardiotoxicity through breast cancer patient specific iPSC-derived heart organoid. Heliyon 2024;10(20); doi: 10.1016/j.heliyon.2024.e38714.

24. Richards DJ, Li Y, Kerr CM, et al. Human cardiac organoids for the modelling of myocardial infarction and drug cardiotoxicity. Nat Biomed Eng 2020;4(4):446–462; doi: 10.1038/s41551-020-0539-4.

25. Moshksayan K, Harihara A, Mondal S, et al. OrganoidChip facilitates hydrogel-free immobilization for fast and blur-free imaging of organoids. Sci Rep 2023;13(1); doi: 10.1038/s41598-023-38212-8.

26. Moshksayan K, Harihara A, Mondal S, et al. A Microfluidic Chip with Immobilization Chambers for Cardiac Organoid Imaging. SPIE-Intl Soc Optical Eng; 2023; p. 8; doi: 10.1117/12.2647888.

27. Zhou Y, Richards AM, Wang P. Characterization and Standardization of Cultured Cardiac Fibroblasts for Ex Vivo Models of Heart Fibrosis and Heart Ischemia. Tissue Eng Part C Methods 2017;23(7):422–433; doi: 10.1089/ten.tec.2017.0169.

28. Li X, Zhang Y, Jin Q, et al. Silicate Ions Derived from Calcium Silicate Extract Decelerate Ang II-Induced Cardiac Remodeling. Tissue Eng Regen Med 2023;20(5):671–681; doi: 10.1007/s13770-023-00523-2.

29. Kumar S, Singh P, Kumar A. Targeted Therapy of Irritable Bowel Syndrome with Anti-Inflammatory Cytokines. Clin J Gastroenterol 2022;15(1); doi: 10.1007/s12328-021-01555-8.

30. Patel P, Karch J. Regulation of Cell Death in the Cardiovascular System. In: International Review of Cell and Molecular Biology Elsevier Inc.; 2020; pp. 153–209; doi: 10.1016/bs.ircmb.2019.11.005.

