## Supplemental Figures for "Optimizing cardiac organoid culture to enhance maturation, viability, and cardiotoxicity assessments"

### Optimizing cardiac organoid culturing conditions for enhanced maturation, viability, and cardiotoxicity assessments

---

1 - Department of Mechanical Engineering, The University of Texas at Austin, Austin, Texas.

2 - Department of Biomedical Engineering, The University of Texas at Austin, Austin, Texas.

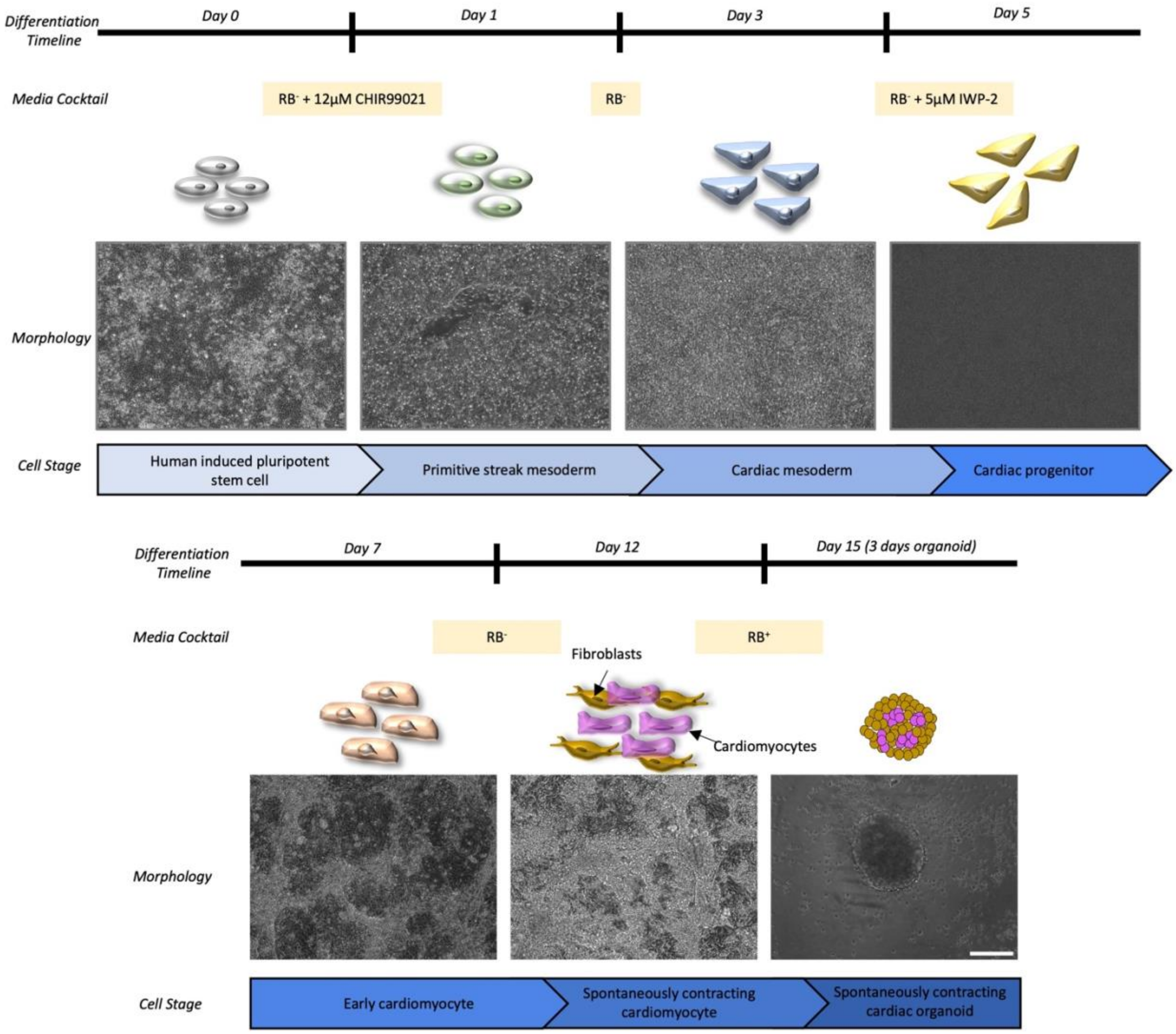


**Supplementary Figure S1**: **Differentiation schematic with representative images of cardiac differentiation.** Differentiation schematic of hiPSC-derived cardiac populations. WNT pathway modulation gives rise to cardiac cells comprising primarily of cardiomyocytes and cardiac fibroblasts. Widespread spontaneous beating activity is observed after 12 days of 2D differentiation, at which point cells are harvested and made into organoids for optimal organoid determination. Scalebar is 200 µm.

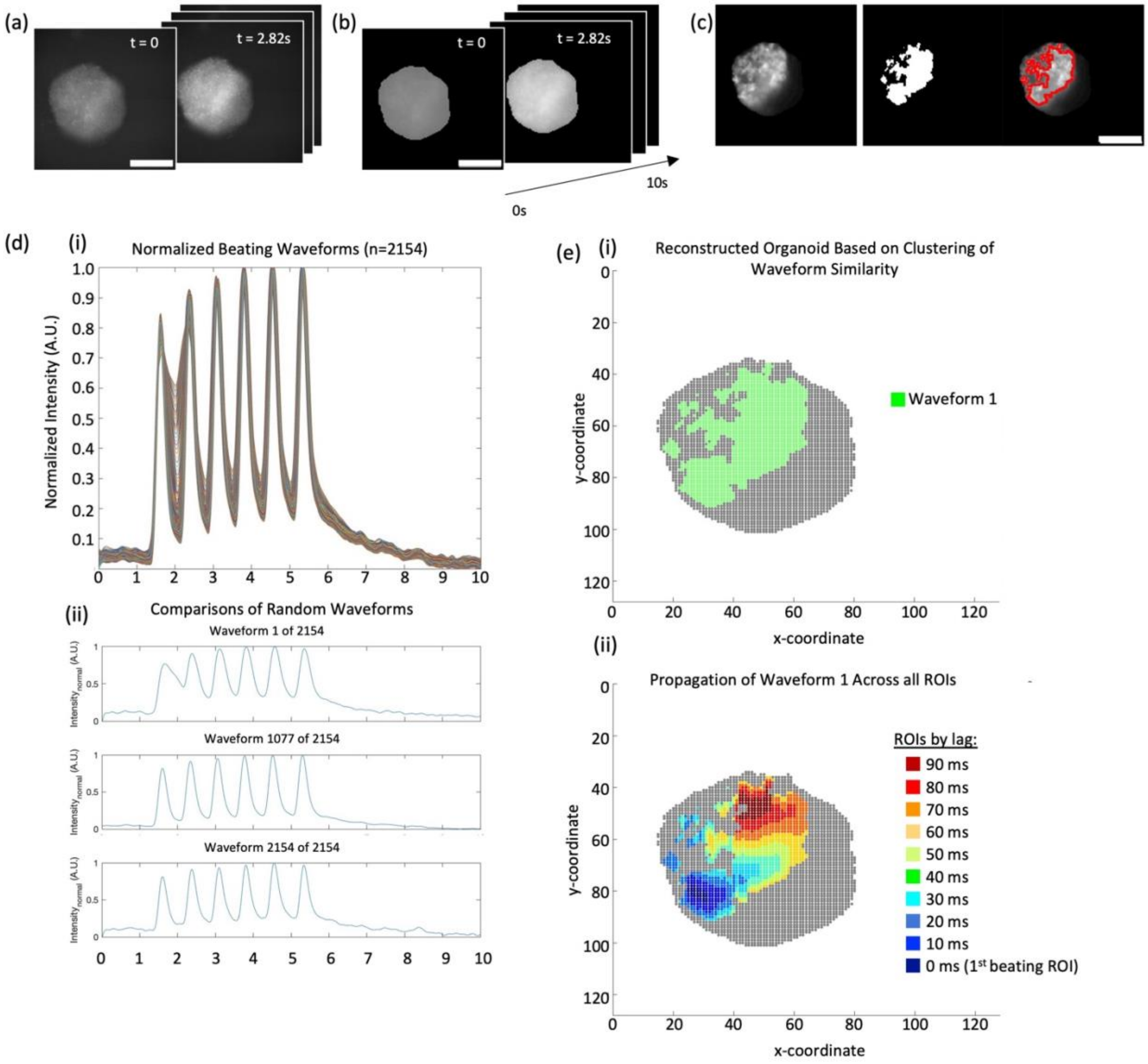


**Supplementary Figure S2**: **Pipeline for Beating Analysis. (a)** Image sequences of $Ca^{2+}$ transients were captured over 10 seconds (scalebar is 400 µm). **(b)** Image frames within the sequence were downsampled by using an 8×8 averaging kernel and organoids within the image sequences were identified using Otsu's segmentation methods to generate the mask of the organoid (scalebar is 400 µm). **(c)** To discern the beating regions within the organoid (*left*) (i.e., those that exhibited $Ca^{2+}$ transients), we evaluated the standard deviation of each kernel within the organoid across all frames in the time sequence and generated an organoid map comprising of standard deviations of GCaMP intensity over time (*middle*). By applying the k-means (k = 2) segmentation algorithm to this projection, we identified the organoids' beating regions, outlined in red (*right*) (scalebar: 400µm). **(d, i)** Plots of normalized waveforms generated by examining all $Ca^{2+}$ transients across 10 s of timelapse imaging on all 8×8 kernels identified as beating were generated, with plots of singular, randomly selected waveforms shown (d, ii). **(e, i)** Beating kernels were then cross correlated with one another to determine the homogeneity of the waveform across the organoid. Using density-based scanning algorithms, we clustered waveforms with high cross correlation coefficients and low distance metrics to ensure that similar waveforms were clustered to assess waveform homogeneity within the organoids, as shown by the reconstructed organoid image of beating kernels clustered based on waveform similarity. **(e, ii)** Lags were measured for each waveform cluster, grouped based on high cross-correlation coefficients, to determine the signal propagation of a waveform across the organoid to report on the peak arrival times of the waveform, as shown by the reconstructed organoid image identifying *peak arrival time* within an identified waveform, i.e., the propagation duration of the calcium waveform across the organoid.

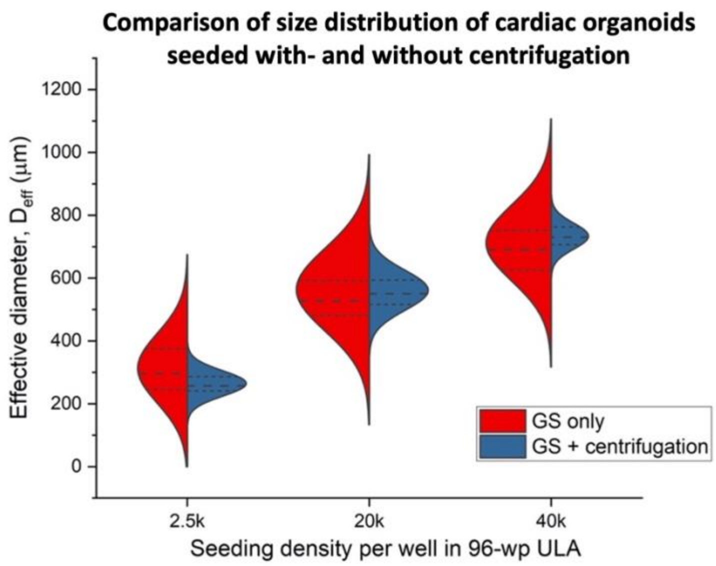


**Supplementary Figure S3: Determination of organoid size distributions upon implementation of a centrifugation step (CS) during organoid formation in 96-well plates.** These data show that the centrifugation step facilitated the generation of more uniformly sized organoids.

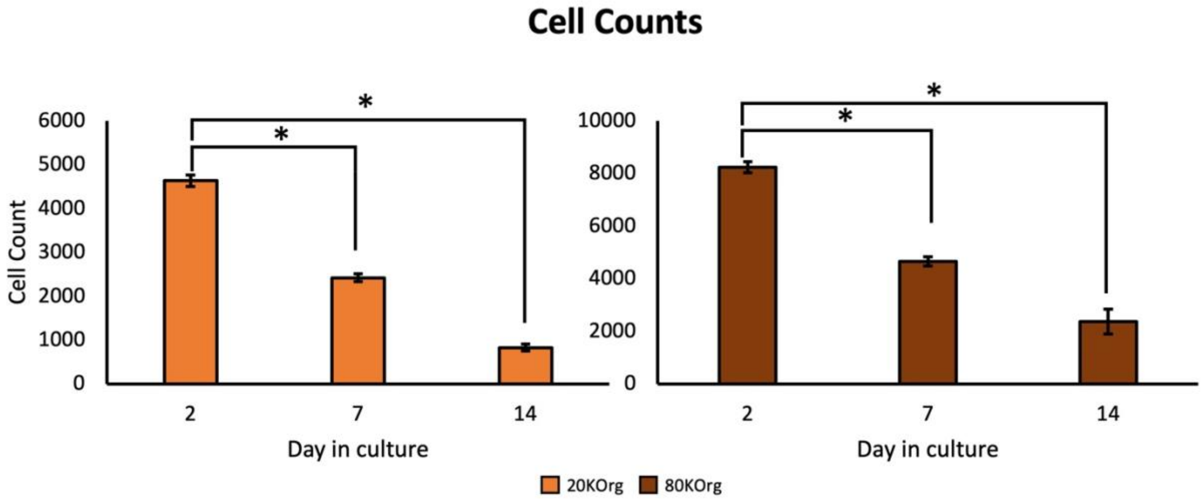


**Supplementary Figure S4: Cell counts of cardiac organoids over time.** DAPI-stained counts of nuclei generated from immunostained cleaved caspase 3 images. Counts were made via 3D imaging volumes of initial cell layers using confocal microscopy. There is significantly less $DAPI^{+}$ nuclei on images counted on Days 7 and 14, when compared to Day 2 for 20KOrg and 80KOrg ($n_{20KOrg, day2}$=4, $n_{80KOrg, day2}$=9, $n_{20KOrg, day7}$=4, $n_{80KOrg, day7}$=5, $n_{20KOrg, day14}$=7, $n_{80KOrg, day14}$=5).

(a)

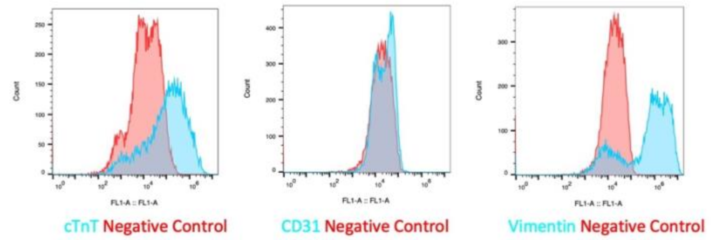


(b)

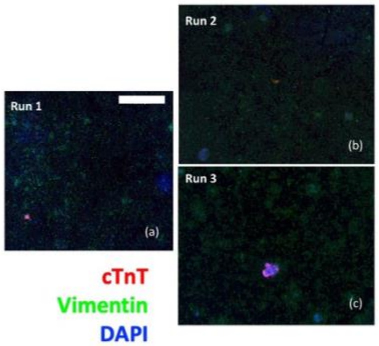


(c)

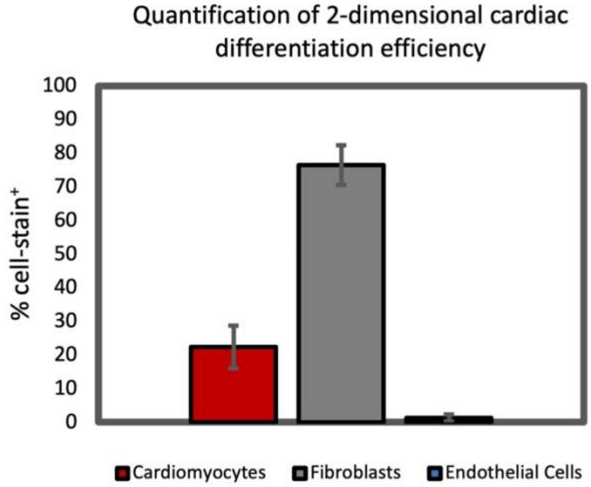


| Cell type | Stain | Percentage |
|---|---|---|
| Cardiomyocyte | cTnT | 22.31 ± 6.38% |
| Fibroblast | Vimentin | 76.40 ± 5.88 % |
| Endothelial Cells | CD31 | 1.29 ± 1.01 % |

**Supplementary Figure S5**: **Protein expression of 2D cells and early organoid cultures probed via flow cytometry.** (**a**) Sample gating strategies applied for quantifying cell-specific antibody staining. Fluorescence histogram overlays of negative controls on experimental samples for determination of cell-specific antibody detection. (**b**) Immunostained cells for cTnT (red) and Vimentin (green), co-stained with DAPI (blue) on cells plated down upon harvesting for organoid formation. Scale bar is 1 mm. (**c**) Summary of 2-dimensional cardiac efficiency quantifying cardiomyocyte, fibroblast, and endothelial cell populations in starting material used for organoid formation. Bar graph demonstrates that key cell types comprising 2D differentiations and cell and organoids at Day 2 of differentiation are cTnT-positive cardiomyocytes and vimentin-positive fibroblasts. Obtained composition data advised gene expression qPCR panel to probe maturity of cardiomyocytes as well as fibrotic activity of fibroblasts (see Figure 2). Table contains representative summary of one characterization of differentiation efficiency (n=6).

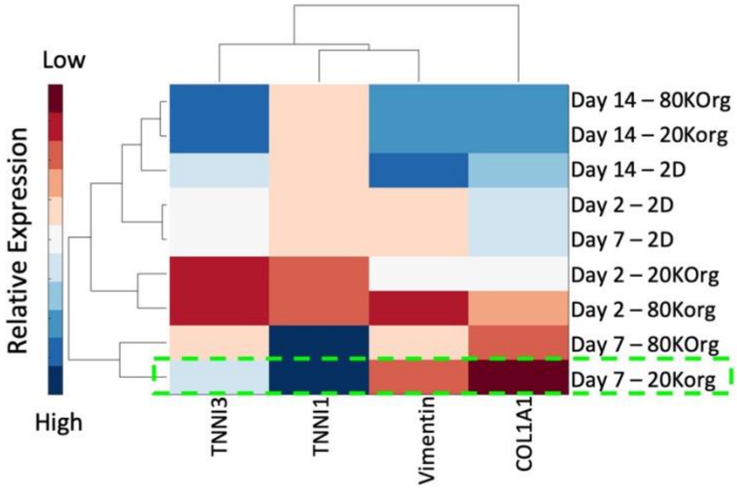


**Supplementary Figure 6:** Clustergram showing unsupervised clustering of 2D and organoid samples based on average $\Delta\Delta C_T$ values of samples.

**Supplementary Table S1:** Complete Primer List Utilized for qPCR

| Gene | Primer Sequence | | Manufacturer |
|---|---|---|---|
| TBP | Forward | 5’-GCTGTTTAACTTCGCTTCCG-3’ | Integrated DNA Technologies |
| | Reverse | 5’-CAGCAACTTCCTCAATTCCTG-3’ | |
| TNNI1 | Forward | 5’-GAACAAGGTGCTGTCTCACT-3’ | Integrated DNA Technologies |
| | Reverse | 5’-CCAGCATTCCTTGGCCTT-3’ | |
| TNNI3 | Forward | 5’-CAGGACTTGTGCCGACAG-3’ | Integrated DNA Technologies |
| | Reverse | 5’-CGCTTAAACTTGCCTCGAAG-3’ | |
| VIMENTIN | Forward | 5’-TGTCCAAATCGATGTGGATGTTTC-3’ | Integrated DNA Technologies |
| | Reverse | 5’-TTGTACCATTCTTCTGCCTCCTG-3’ | |
| COL1A1 | Forward | 5’-GATTCCCTGGACCTAAAGGTGC-3’ | Bio-Rad |
| | Reverse | 5’-AGCCTCTCCATCTTTGCCAGCA-3’ | |
